# Long-term effects of early-life thermal fluctuations on the cellular stress response and CT_max_ of zebrafish, *Danio rerio*

**DOI:** 10.64898/2026.06.03.729897

**Authors:** Hossein Haghighi, Benjamin W. Lindsey, Kenneth M. Jeffries

**Affiliations:** Department of Biological Sciences, University of Manitoba, Winnipeg, Canada; Department of Human Anatomy and Cell Science, Rady Faculty of Health Sciences, University of Manitoba, Winnipeg, Canada

**Keywords:** gene expression, heat shock proteins, oxidative stress, temperature increase

## Abstract

Temperatures in aquatic ecosystems have been affected by anthropogenic activities such as agricultural and industrial water use, and climate change caused by greenhouse gas emissions. Changes in water temperature directly affect the cellular and organismal physiology of fishes because most fishes are ectotherms. These effects can take different forms, such as increased cellular stress and reactive oxygen species (ROS) production. However, the type and magnitude of response depend on the duration and the frequency of the exposure to elevated water temperature. In this study, we investigated the long-term effects of exposure to daily thermal fluctuations occurring during early-life stages of zebrafish, *Danio rerio*, on gene expression and CT_max_ in later developmental stages. To do so, wild-type zebrafish were exposed daily to a + 5°C fluctuation in temperature from ambient (28 °C) to 33 °C over the first 30 days post fertilization (dpf), before being held until 90 dpf at ambient temperature. The fish that experienced daily thermal fluctuation were compared to a control group that was kept at 28 °C throughout the experiment and sampled at the same timepoints. Samples were collected at 18 (larval), 30, 60 (juvenile), and 90 (adult) dpf to study the expression of heat shock proteins and oxidative stress genes. The thermotolerance of fish was tested using CT_max_ trials at 60 and 90 dpf. Daily thermal fluctuation over the first 30 dpf led to a significant increase in the expression of *hsp47*, *gstp1a*, *sod1*, and *sod2* genes at 60 dpf, and *hsp47*, *hsp90aa1*, *hsp90ab1*, *cat*, *glulb*, *gstp1a*, *sod1*, and *sod2* genes at 90 dpf. The only significant increase detected during the larval stage was *glulb* at 18 dpf. Fish that experienced thermal fluctuation also had a higher CT_max_ at 60 dpf, but this increased thermotolerance significantly decreased from 60 to 90 dpf, where it was not different between treatments. Overall, our study demonstrates that early-life thermal stress increased cellular stress responses and thermotolerance in zebrafish into later ontological stages.

## Introduction

Water temperature in freshwater ecosystems often fluctuates on diurnal, seasonal, and annual time scales (Komatsu, 1985; Zhang et al., 2016; Żelazny et al., 2018). Aquatic ectotherms have adapted to tolerate natural fluctuations in water temperature (Goldspink, 1995). However, the temperature of aquatic ecosystems has been influenced by human activities (McBryan et al., 2013). For example, the discharge of warm water from power plants and agricultural and farming activities have altered the natural thermal regime of aquatic ecosystems, causing fluctuations in water temperature (Bukola et al., 2015; Guimarães, 2023). Further, climate change has also affected the magnitude and intensity of heatwaves in aquatic ecosystems (Woolway et al., 2021; Woolway et al., 2022). High temperatures can directly affect fish populations and, in extreme cases, can result in mortality (Ern et al., 2023).

Most fishes are ectotherms and environmental temperature can directly affect their physiology, metabolism, and behavior (Alfonso et al., 2020). Thermal stress has been reported to trigger an organismal stress response through increased cortisol and glucose levels in the blood (Kim et al., 2019). At the cellular level, the upregulation of genes such as heat shock proteins (HSPs), which code proteins that function as protein chaperones, has also been observed in numerous different species (Komoroske et al., 2015; Jeffries et al., 2018; Kim et al., 2019; Bugg et al., 2020; Mackey et al., 2021; Gavarikar and Craig, 2025). Additionally, thermal stress has also been reported to increase oxidative stress in fishes, which causes oxidation in lipids and other structural molecules (Lushchak and Bagnyukova, 2006). This increase in oxidation of biomolecules can result in the upregulation of the expression of antioxidant proteins that are involved in an oxidative stress response (Madeira et al., 2013). Increased temperature can also affect the susceptibility of fish to xenobiotics and reduce the immune system response to pathogens (Dietrich et al., 2014; Bugg et al., 2023). Increased upregulation of genes coding for apoptosis and proinflammatory proteins, such as IL-7, IL-8, IL-10, and IL-1β, has also been reported following thermal shock (Liu et al., 2022). Collectively, changes in temperature affect the cellular stress response in fish, which can reduce their performance and potentially affect fitness (Jeffries et al., 2018).

In fishes, thermal stress during the early larval life stages can impact metabolism, growth rate, and survival (Pankhurst and Munday, 2011). For instance, increasing temperature in the embryonic stage has been shown to increase the number of abnormal embryos in *Danio rerio*, decrease hatching success in *Trichogaster leeri*, and accelerate the beginning of endogenous feeding in *Symphysodon aequifasciatus* (Schirone and Gross, 1968; Pereira et al., 2016; Mattos et al., 2024). ٍMoreover, exposure to increased temperature in the larval stage has been shown to result in developmental deformities in *Labeo rohita*, changes in body composition in *Inimicus japonicus*, increased activity of larvae in *Paralichtys olivacous*, and increases in the pelagic larval period and swimming ability in *Amphiprion melanopus* (Fukuhara, 1990; Green and Fisher, 2004; Wen et al., 2013; Ashaf-Ud-Doulah et al., 2021). Overall, exposure to temperatures higher or lower than the species-specific ideal range can have adverse effects on growth in fish in different life stages and increase morphological abnormalities and mortality (Steinarsson and Björnsson, 1999; Abdel et al., 2004; Le et al., 2011; Tsuji et al., 2014; Shahjahan et al., 2018; Islam et al., 2020). Additionally, temperature strongly influences the developmental rate of fish during early life stages (Kimmel et al., 1995), and their limited thermal tolerance at this sensitive developmental window makes them especially vulnerable to fluctuations in temperature (Dahlke et al., 2025).

Early-life exposure to stress has been reported to have long-lasting effects on animals (Lux, 2018). Although some of these effects can be beneficial and improve animals’ resilience to subsequent stressors, they can also have detrimental effects (Cramer et al., 2019). For instance, non-lethal thermal stress during early life has been reported to cause long-lasting epigenetic changes and increase the oxidative stress response in adult animals (Costantini et al., 2012; Zhou et al., 2018). The long-term effects of constant thermal conditions have been studied in several species; however, literature on the long-term effects of fluctuating temperatures remains limited (Blanchard et al., 2025). Because thermal stress has been reported to affect the physiology of animals from different perspectives such as oxidative stress response, and energy use, and protein metabolism (Kumar et al., 2015; Madeira et al., 2016; Resende et al., 2022; Khieokhajonkhet et al., 2023), studying the possible long-term effects of temperature on the cellular stress response and thermotolerance across different life stages in fishes can increase our understanding of thermal plasticity in animals.

In this study, we investigated the long-term effects of exposure to daily thermal fluctuation at the embryonic and larval stages of zebrafish. We hypothesized that experiencing early-life thermal fluctuations affects the expression of genes involved in general stress and oxidative stress responses and also modifies thermotolerance in fish. To test this hypothesis, we exposed zebrafish to daily thermal fluctuations during the embryonic to early larval stage (0-30 days post-fertilization; dpf), and by using qPCR, we measured the mRNA transcript abundance of genes involved in a cellular stress response from the larval through to the adult stage (90 dpf). To determine if daily thermal fluctuations affect the thermotolerance of fish in later life stages, Critical Thermal Maximum (CT_max_) trials were further conducted on juvenile and adult fish at 60 and 90 dpf, respectively. Additionally, we employed transcriptomic markers to estimate the sex of individuals based on gene expression in order to analyze relative gene expression for males and females separately at 60 and 90 dpf. This work contributes to a growing number of studies that demonstrate how different thermal treatments during early life stages can lead to delayed effects at later life stages, affecting growth, thermotolerance, and stress responses in fish.

## Materials and Methods

### Fish breeding and husbandry

Wild-type AB (WT-AB) zebrafish were bred and reared in the Rady Biomedical Fish Facility at the University of Manitoba, Winnipeg, Manitoba, Canada, on a 12 light (7 am):12 dark (7 pm) light cycle unless stated otherwise. Fish were housed on a recirculating Tecniplast housing system (Techniplast Inc. USA), equipped with automatic dosing, UV and carbon filters, with a stable pH and conductivity of 7.4 and 900 µS, respectively. Facility water was maintained at 28°C.

For breeding, two females and three males were introduced together in the evening into a zebrafish breeding tank with a meshed bottom to separate the fish from the eggs and facilitate embryo collection the following day. To mimic the natural habitat, artificial plants were placed in the tank. Breeding occurred the following morning with initiation of the light cycle, stimulated by the gradual increase in light intensity. Fertilized eggs were collected and transferred to Petri dishes with embryo medium (0.01 mg.L^-1^ methylene blue dissolved in fish facility water) at a maximum density of 25 embryos dish^-1^. Petri dishes were then transferred to a Fisherbrand 18L Low Temp Incubator (Thermo Fisher Scientific, Pittsburgh, USA). To reduce the potential for vertical transmission of pathogens and fungi from parents to offspring, embryos were bleached at 1 dpf by submerging in 7 mg.L^-1^ chlorine for 5 min, followed by three washes with embryo medium, each for 3 min in three separate containers. Incubation of eggs in the incubator continued until 5 dpf, when almost all the eggs were hatched.

Upon depletion of the yolk sac at 5dpf, larvae were fed two portions of GEMMA Micro 75 (Skretting, Maine, USA) and one portion of live rotifer daily. Juvenile fish (> 30 dpf) were fed two portions of GEMMA Micro 150 and one portion of brine shrimp nauplii (Brine Shrimp Direct, Ogden, USA) daily. Temperature, oxygen, and ammonia in the tanks were measured throughout the experiment.

### Temperature treatment

Immediately following embryo collection at 0 dpf, zebrafish were separated into two groups: (1) temperature treatment, and (2) control. From 0-6 dpf, the fish were held in the Low Temp incubator, where the temperature regimes were maintained for the two groups. At 6 dpf, larvae were transferred to 18 (L) × 14 (W) × 10 (H) cm aerated containers filled with fish facility water up to 3 cm until 30 dpf. Then, the containers were placed in 56 (L) × 30 (W) × 30 (H) cm coolers. The coolers were filled with distilled water, and the temperature was maintained at 28 °C by using electric heaters placed at the bottom of each cooler. To maintain homogeneous temperature in the coolers and subsequently in the containers, the water in the coolers was constantly mixed with a pump. Fish were kept in these containers until the end of the larval stage (30 dpf), after which fish were transferred to standard 3.5 L Tecniplast Inc. zebrafish tanks (27 (L) × 11 (W) × 16 (H) cm) and reared on a Tecniplast recirculating aquatic housing system at 28 °C until the end of the experiment (90 dpf) as described above.

In the temperature treatment group, at 12:00 pm daily, prior to feeding, the temperature was raised to 33 °C from ambient 28 °C water at the rate of ∼0.8 °C min^-1^ and then decreased at the same rate back to 28 °C over a 2-hour period during the embryonic and larval stages (0-30 dpf). The control group experienced similar handling but did not experience any thermal fluctuations during embryonic and larval stages, and the temperature for this group was kept constant at 28 °C throughout. In addition, the total length at 15 and 30 dpf, and fork length at 60 and 90 dpf were measured and compared between the two groups using the Mann-Whitney U test (p-value < 0.05).

### Critical Thermal Maximum (CT_max_) trials

Acclimation to higher temperatures has been shown to increase the CT_max_ in fishes (Bugg et al., 2020; Waterbury et al., 2024). To test if daily thermal fluctuations can have effects on the CT_max_ in fishes, we conducted CT_max_ on 60 and 90 dpf zebrafish that experienced daily thermal fluctuations during embryonic and larval stages and compared it with the control group. The trials were conducted in an 18 (L) × 14 (W) × 10 (H) cm container filled with fish facility water up to 8 centimeters. The containers were placed in coolers filled with distilled water, and the temperature was maintained at 28 °C. Three hours before the CT_max_ trials, nine to fifteen fish were introduced into separate well-aerated containers. To conduct the CT_max_, we increased the temperature in the containers at a target rate of 0.3 °C min^-1^ using an electric heater in the cooler. The temperature was monitored using digital thermometers and recorded using temperature probes (Witrox Oxygen probe, Loligo Systems; Viborg, Denmark). The behavior of the fish was monitored constantly until they lost equilibrium and could not re-orientate themselves. When fish lost equilibrium and could no longer re-orient, the temperature was recorded, and the fish was netted and euthanized in a separate tank using tricaine (0.4%, pH 7.4). A two-way ANOVA followed by a Bonferroni correction was used to test the significant difference (p-value < 0.008) between groups.

### Relative gene expression

We collected 10 fish from each group at 18, 30, 60, and 90 dpf (10 individuals × 2 groups × 4 timepoints) to study changes in gene expression. Samples were collected in the morning, and after euthanizing with tricaine (0.4%, pH 7.4), samples were preserved in RNA*later* (Thermo Fisher Scientific, Waltham, USA). After 24 h of preservation at 4 °C, samples were transferred to a -80 °C freezer for storage. Total RNA was extracted using MagMax mirVana™ Total RNA Isolation Kit (Applied Biosystem, Germany) on a KingFisher Duo Prime (ThermoFisher Scientific, USA) machine. Briefly, the whole fish was homogenized using a TissueLyser (QIAGEN, Germany) at 50 Hz for 5 min in the lysis buffer provided by the kit. The homogenized tissue was centrifuged and transferred to an extraction plate, and RNA purification was conducted following the manufacturer’s protocol. The quantity and purity of extracted RNA were measured using a NanoDrop One spectrophotometer (ThermoFisher Scientific). For RNA integrity, we loaded 400 ng of RNA on a 1% agarose gel and checked the 18S and 28S rRNA bands. Before cDNA synthesis, 300 ng of total RNA from each sample was treated with Invitrogen™ *ezDNase* enzyme (Thermo Fisher Scientific, USA) at 37 °C for 2 min to eliminate genomic DNA. Then, the DNase-treated RNA was used for cDNA synthesis using a High-Capacity cDNA Reverse Transcription Kit (ThermoFisher Scientific) following the manufacturer’s instructions.

We used qPCR to measure and compare the expression of four genes involved in cellular heat shock response and seven oxidative stress response genes (**Table 1**). Sequences of genes were downloaded from the National Center for Biotechnology Information (NCBI) (www.ncbi.nlm.nih.gov/), and primers were designed using Primer Express (3.0.1) (Thermo Fisher Scientific). For qPCR, the synthesized cDNA was diluted 1:4 with nuclease-free water. Then, 5 µl of the diluted cDNA was mixed with 6 µl of PowerUp™ SYBR™ Green Master Mix (ThermoFisher Scientific, USA), 0.1 µl of forward primer (10 mM), 0.1 µl of reverse primer (10 mM), and 0.8 µl of nuclease-free water in a 384-well plate. The qPCR was performed using a QuantStudio 5 (Applied Biosystems, USA) machine, and the thermal conditions for qPCR included initial denaturation at 95 °C for 5 min, 40 cycles of denaturation at 95 °C for 15 s, annealing at 58 °C for 15 s, and extension at 72 °C for 20 s. Melt-curve analysis was also included in the analysis to confirm that there was no non-specific amplification or primer-dimer formation. The efficiency of primers was measured by generating standard curves using serial dilution. The method described by Pfaffl (2001) was used to analyze gene expression data using three internal control genes for normalization. Because the distribution of data was not normal as determined by Shapiro–Wilks’s tests, a Mann-Whitney U test was used to determine significant differences (p-value < 0.05) between the temperature treatment and control group. The same test was also used to compare the difference in the expression of each gene between males and females.

**Table 1.**
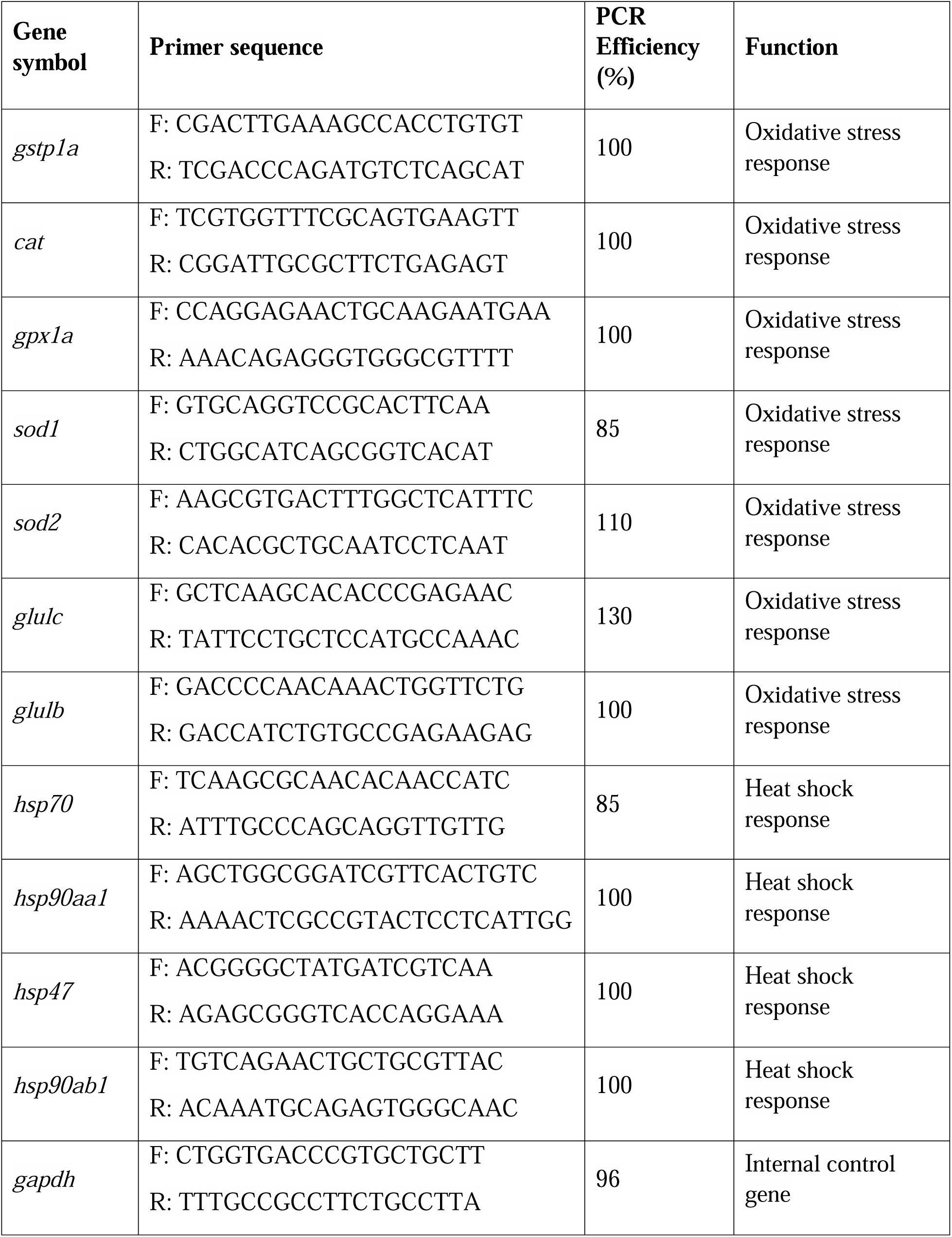
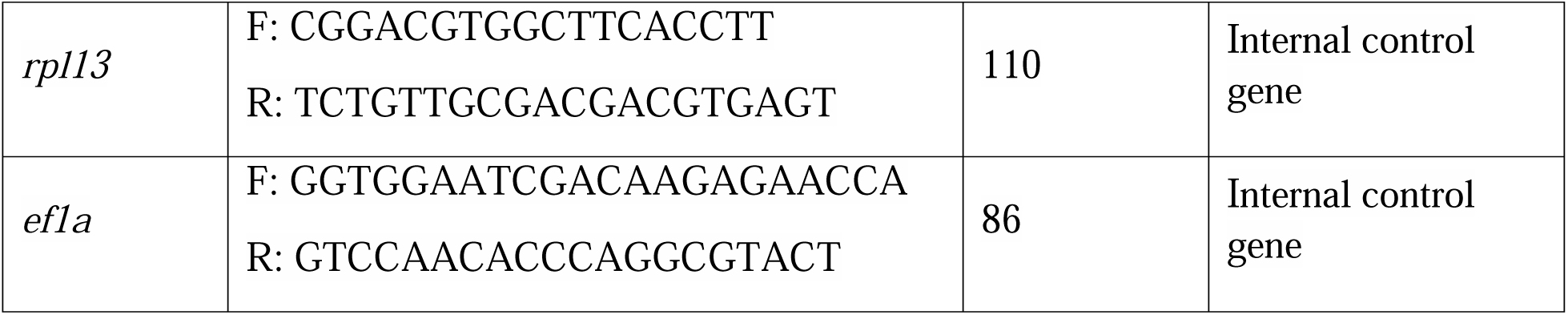
List of primers used for amplification of oxidative stress and heat shock response genes in zebrafish, *Danio rerio*, along with internal control genes.

### Sex determination

We attempted to determine the sex of each fish before gene expression analysis to avoid misinterpretation of results caused by sex-biased expression of oxidative stress and HSP response detected in fish species (Li et al., 2014; Błońska et al., 2021). Due to difficulties in distinguishing males from females through visual inspection, especially in younger fish, and also lack of genetic sex markers, we used transcriptomic methods for sex determination. To determine the sex of each fish based on gene expression, twelve genes that were previously reported to have sex-biased expression were chosen (Wong et al., 2014; King et al., 2020; **Table 2**). To test the accuracy of these genes for sex determination, the expression patterns were tested on 5 adult male and 5 adult female zebrafish (10-month-old) with clear external sex-specific characteristics that were visible under a microscope. Since the candidate genes used for the determination of sex may be expressed in different organs, such as the brain and the ovary, RNA was extracted from a homogenized sample of each fish. To do so, the entire fish was frozen in liquid nitrogen and ground in a mortar using a pestle. Total RNA extraction, cDNA synthesis, and qPCR were conducted as before. Then, the expression of these twelve genes in females was measured relative to males using Pfaffl’s method (Pfaffl, 2001). The sequence of primers used for amplifying these genes in qPCR, along with the expression of each gene in females relative to males, is listed in **Table 2**. Among the twelve genes tested, *bmp15*, *ccnb1*, *gyg1a*, *kpna2*, *rdh10b*, and *zp3* showed higher differences in expression between males and females and were chosen to be used in this study.

**Table 2.**
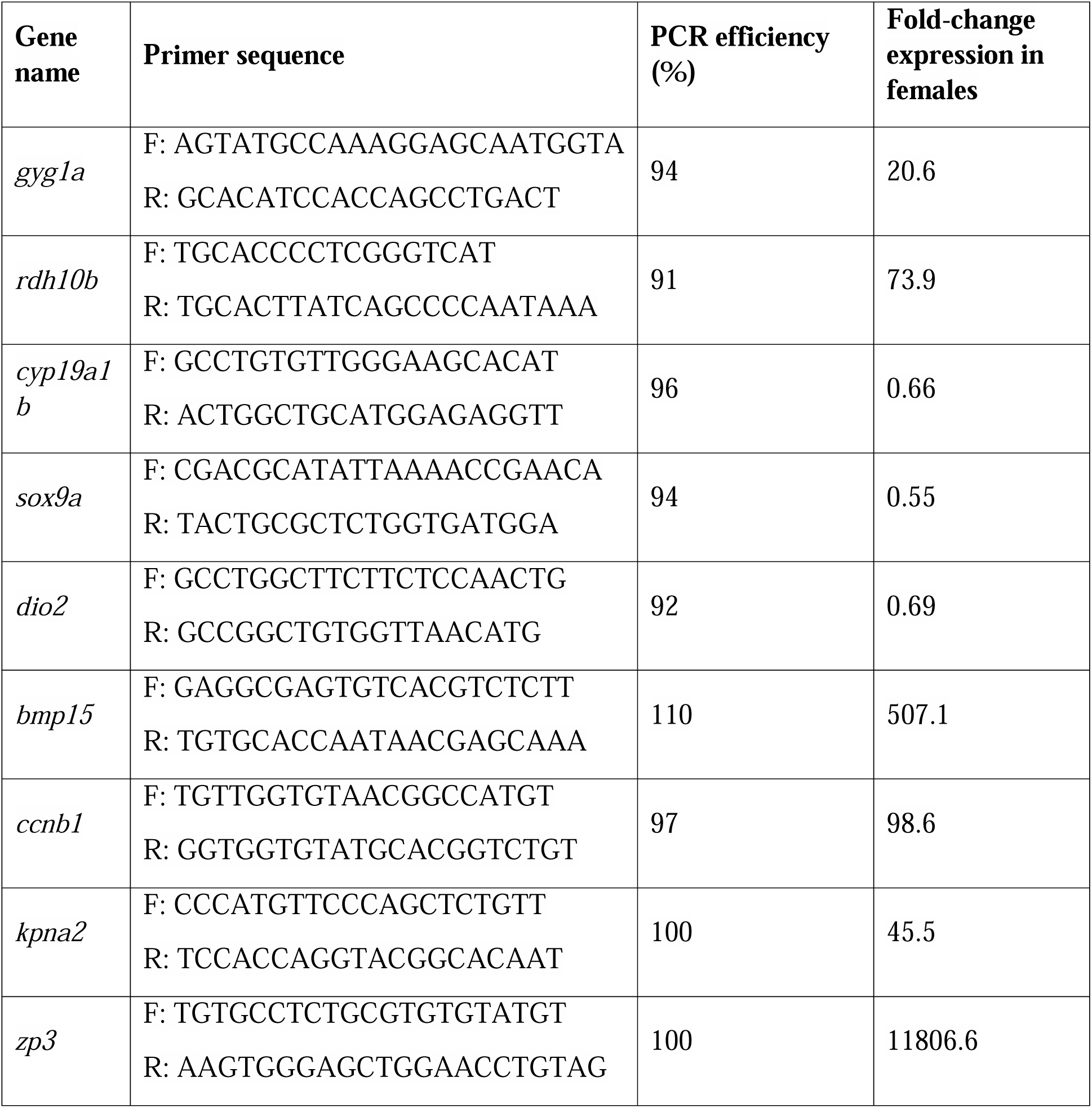
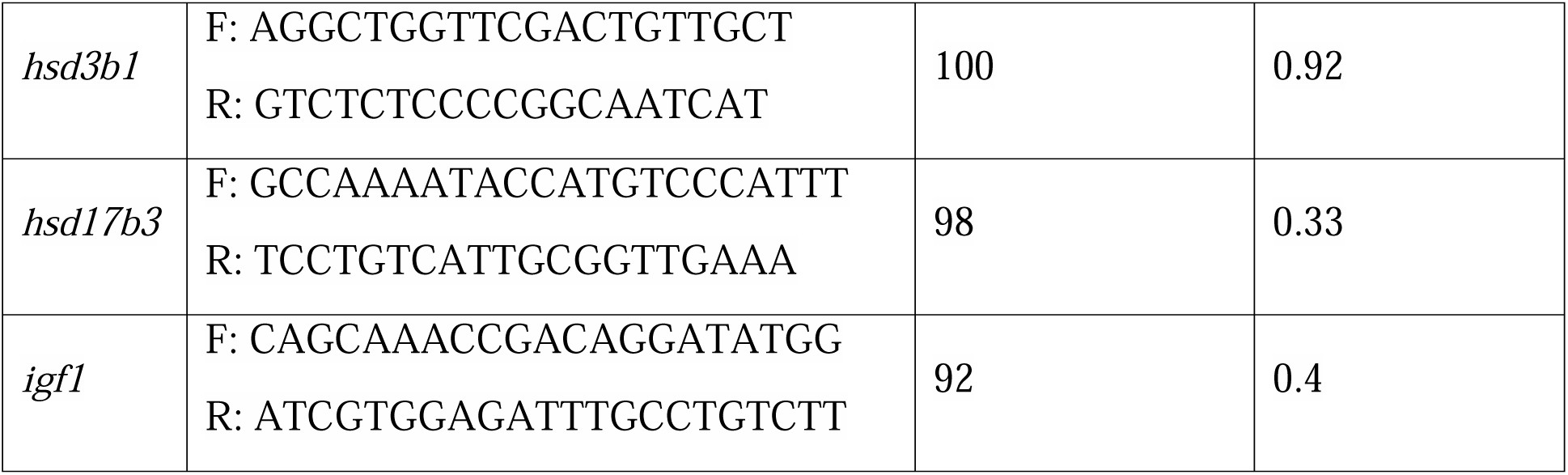
List of primers and probes for amplification of genes used for sex determination in zebrafish, *Danio rerio*, along with their fold-change expression in females relative to males.

We used a Random Forest technique to determine the sex of individuals from the thermal exposure trials using the expression of genes with sexually biased patterns. To do so, the expression of *bmp15*, *ccnb1*, *gyg1a*, *kpna2*, *rdh10b*, and *zp3* was measured for each individual using qPCR, and the C_t_ (cycle of threshold) value was normalized to the geometric mean of internal control genes *ef1a*, *gapdh*, and *rpl13* (Ct _(target gene)_-Ct _(control gene)_). Then, the normalized C_t_ values from adult fish with known sex were used to train a Random Forest model using the Random Forest package in R (Liaw and Wiener, 2002). The performance of the model was evaluated using the classification accuracy test, which is defined as the proportion of correctly predicted instances among the total number of instances. After training the model, the normalized C_t_ of the six sexually biased genes checked in the individuals from the temperature treatment group and control group was used to estimate the sex of that individual using the trained model.

## Results

The temperature treatment group and the control group did not show a significant difference (p-value > 0.05) in length at 15 and 30 dpf; however, fork length was significantly smaller in the temperature treatment group (p-value < 0.05) at 60 and 90 dpf (**Figure 1**). We found that CT_max_ was significantly higher in the temperature treatment group at 60 (p-value < 0.008), but not at 90 dpf (**Figure 2**). There was a 0.62 °C difference in average CT_max_ between the temperature treatment group and control group at 60 dpf, but this difference was decreased to 0.27 °C at 90 dpf. Additionally, the CT_max_ significantly decreased from 60 dpf to 90 dpf by 0.68 °C in the temperature treatment group (p-value < 0.05), but there was no significant difference in CT_max_ between the control groups at 60 and 90 dpf (p-value > 0.008).

**Figure 1.**
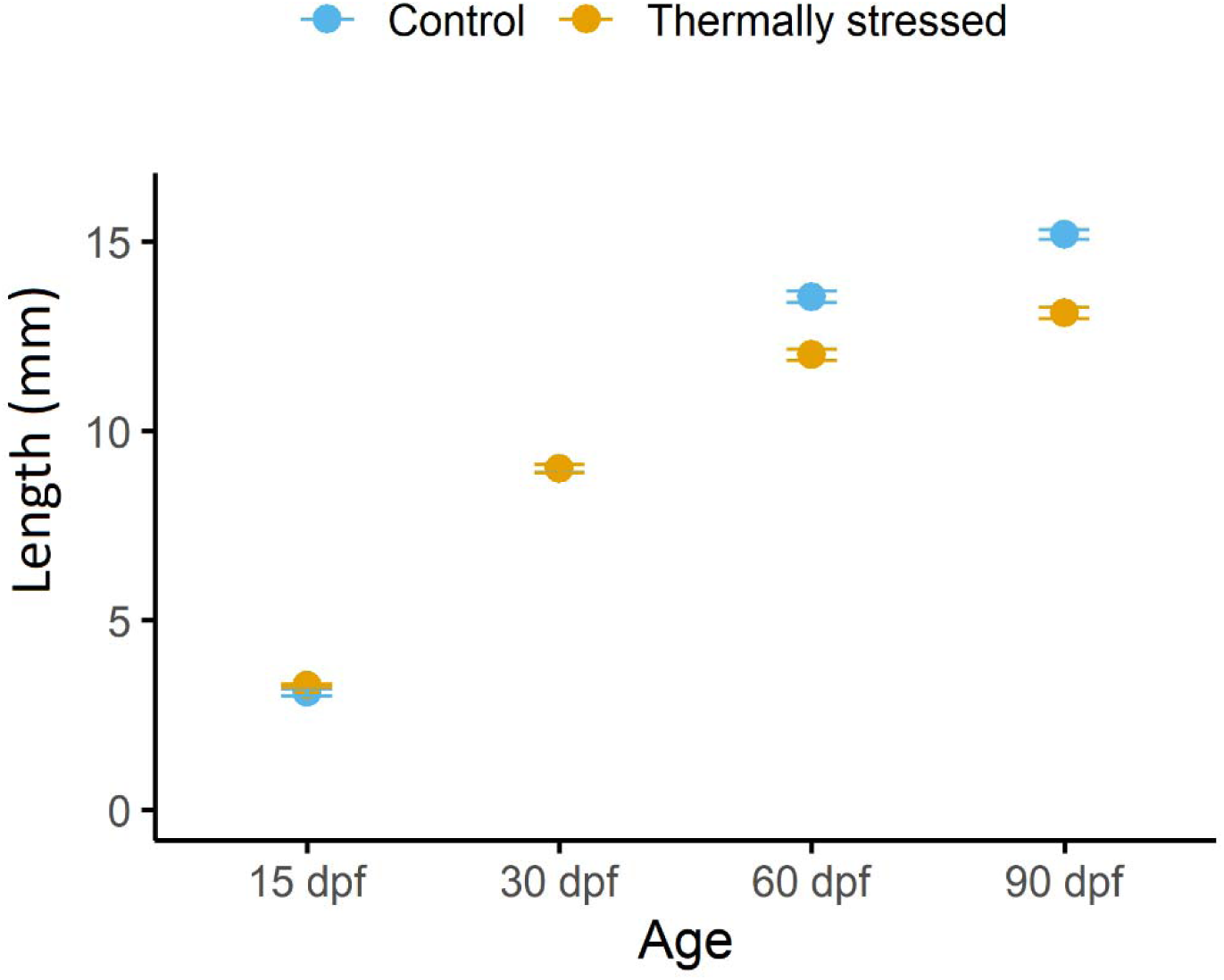
Length of *Danio rerio*, measured at 15, 30, 60, and 90 dpf in the temperature treatment group and control group. Fish in the temperature treatment group experienced daily thermal shock (+ 5 °C) during the larval stage (1-30 dpf) and then were kept at 28 °C until 90 dpf to study the long-term effects of early-life thermal fluctuations. Fish in the control group did not experience any thermal fluctuations and were kept at 28 °C throughout the experiment. At 15 and 30 dpf, total length was measured, but at 60 and 90 dpf, fork length was measured. At 30 dpf, the length data of the control group and temperature treatment group overlap. The statistically significant difference (p-value < 0.05) between groups at each timepoint was measured using Mann-Whitney U tests and shown by using asterisks (*). Error bars show standard error (SE).

**Figure 2.**
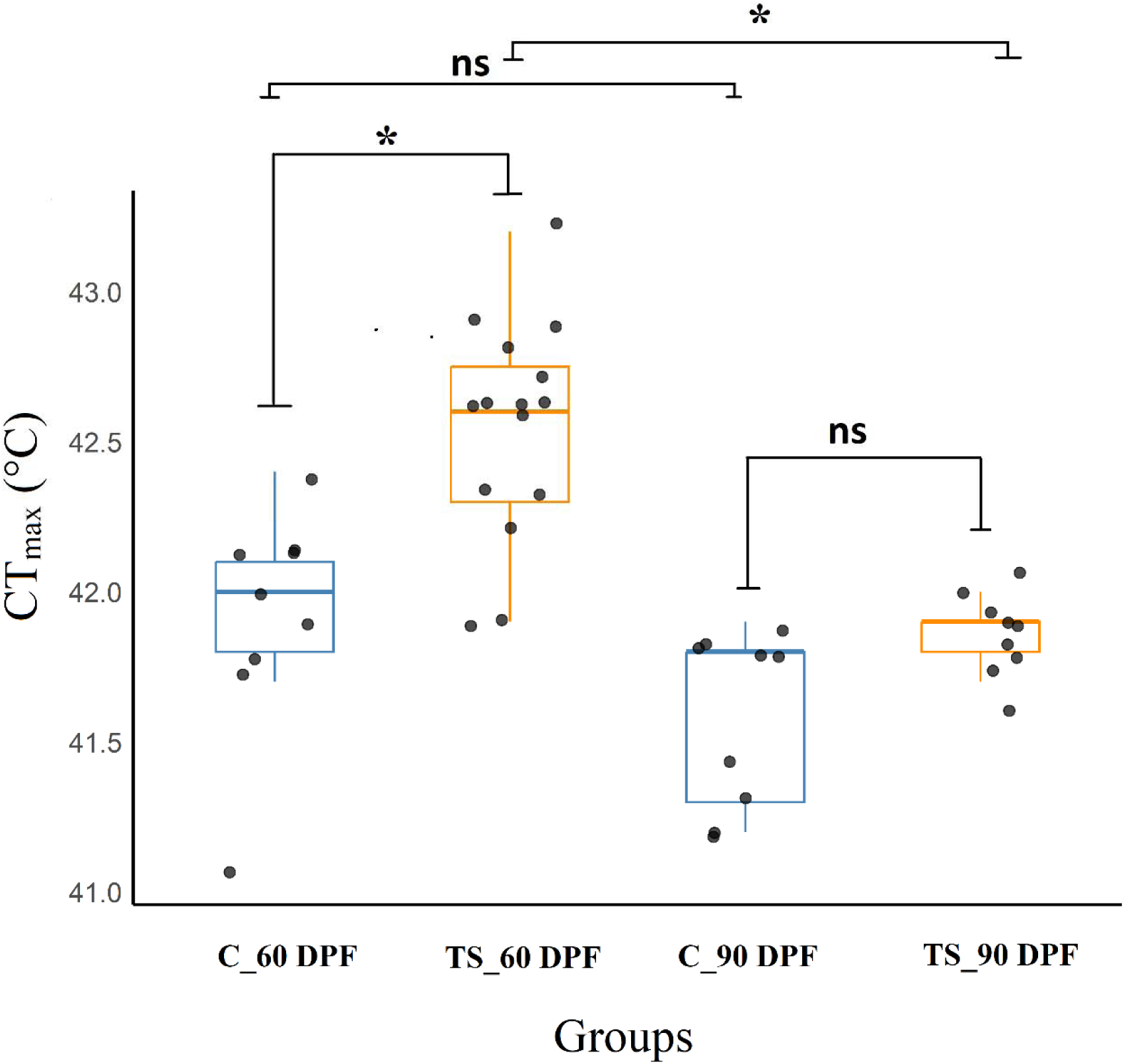
Critical thermal maximum (CT_max_) of *Denio rerio* was measured at 60 and 90 dpf in the temperature treatment (TT) and control (C) group. In the temperature treatment group, fish experience daily thermal fluctuation (+ 5 °C) every day during the larval stage and then are kept at 28 °C during the juvenile stage until the adult stage (90 dpf). The control group was kept at 28 °C throughout the experiment (1-90 dpf) and did not experience any thermal fluctuations. The significant difference (p-value < 0.008) between groups and also between different timepoints tested using the Mann-Whitney U test is shown using asterisks (*).

The classification accuracy test for the Random Forest model showed 100% accuracy of the model in distinguishing males from females using gene expression patterns measured in the fish with known sex. However, when we used the expression of *bmp15*, *ccnb1*, *gyg1a*, *kpna2*, *rdh10b*, and *zp3* to distinguish males from females in the temperature treatment group and control group, it was only feasible at 60 and 90 dpf. At 60 dpf, the sex ratio of males:females was 80:20 in the temperature treatment group, and 67:33 in the control group. At 90 dpf, the sex ratio of males:females was 70:30 in the temperature treatment group, and 40:60 in the control group. Due to a low number of females in the temperature treatment group at 60 dpf, female samples were excluded, and only samples from males were used for gene expression analysis at 60 dpf. However, at 90 dpf, both sexes were included in the analysis.

Although the temperature treatment group did not exhibit a significant change (p-value > 0.05) in the expression of the four *hsp* genes during the larval stage, there was a significant difference (p-value < 0.05) in expression at 60 and 90 dpf (**Figure 3A**). At 60 dpf juvenile animals, *hsp47* was significantly upregulated (2.71-fold) (p-value < 0.05), although the other *hsp* genes did not show a significant change. At 90 dpf, *hsp90aa1* was significantly upregulated (p-value < 0.05) in both males (2-fold) and females (3-fold), and this upregulation was significantly higher (p-value < 0.05) in females compared to males. The expression of *hsp90ab1* was significantly upregulated (p-value < 0.05) in males (1.42-fold) at 90 dpf, while slightly downregulated in females (0.66-fold). Additionally, *hsp47* showed significant (p-value < 0.05) upregulation (2.07-fold) in the females measured at 90 dpf.

**Figure 3.**
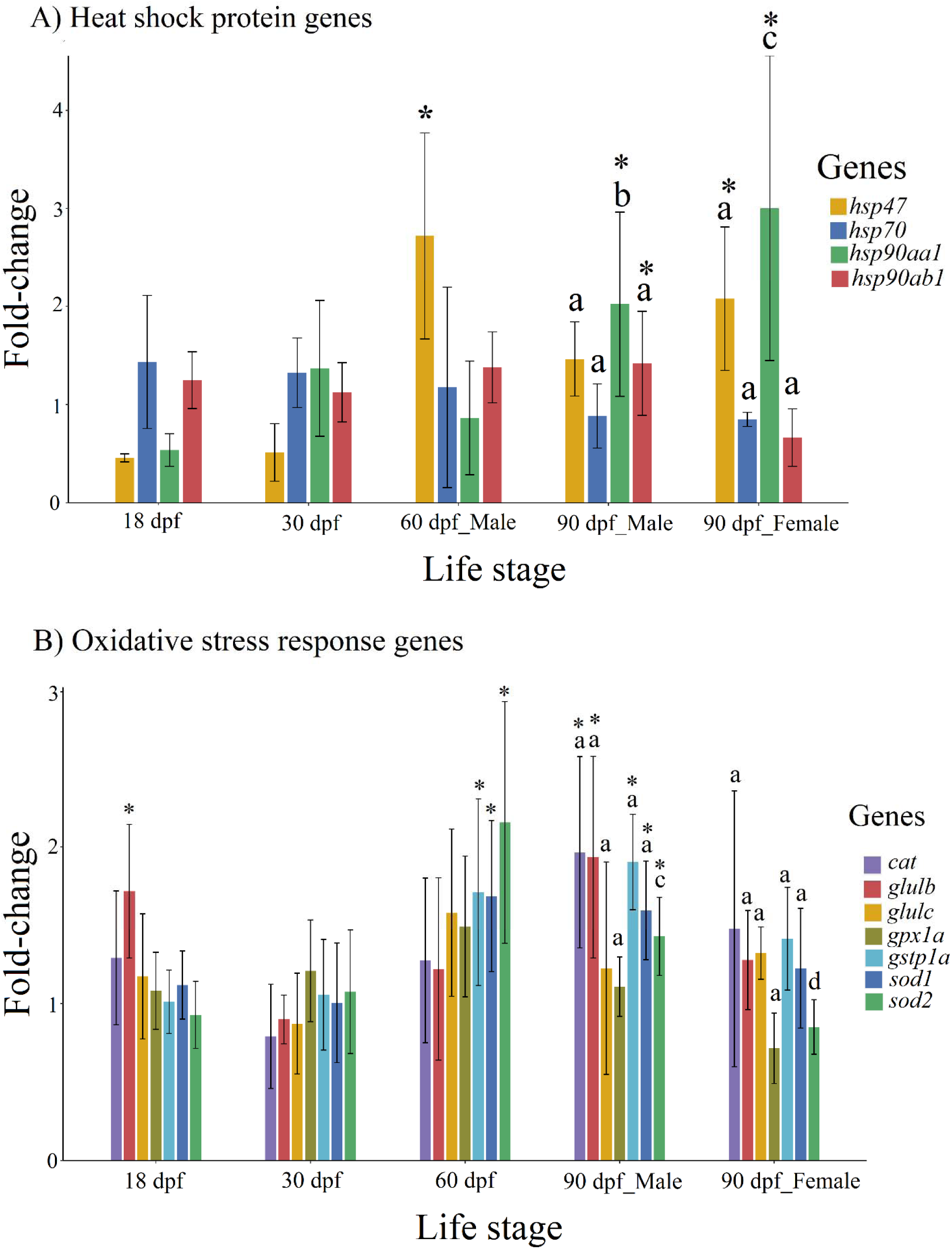
Relative gene expression of genes coding HSPs (A) and proteins involved in oxidative stress response (B) in response to early-life thermal fluctuations in the *Danio rerio*. Fish experienced daily thermal fluctuations (+ 5 °C) during the larval stage (1-30 dpf) and then were kept at 28 °C until 90 dpf to study the long-term effects of early-life thermal fluctuations. The expression of genes at each timepoint was measured compared to a control group that was kept at 28 °C during the experiment. In these graphs, the changes in the expression of each gene are shown as fold-change upregulation or downregulation relative to the control group. The asterisks (*) show the significant differences (P-value < 0.05) between the expression of each gene in the temperature treatment group and the control group at each timepoint. At 60 dpf, only the relative gene expression in the males is shown, and females were omitted from statistical analysis due to a low number. At 90 dpf, relative gene expression was measured separately for males and females, and significant differences (P-value < 0.05) between sexes are shown by different letters for each gene.

During the larval stage, only *glulb* was significantly upregulated (p-value < 0.05) (1.71-fold) at 18 dpf in the temperature treatment group (**Figure 3B**). At 60 dpf, the expression of *gstp1a* (1.49-fold), *sod1* (1.68-fold), and *sod2* (2.15-fold) were significantly upregulated (p-value < 0.05). At 90 dpf, *cat* (1.96-fold), *glulb* (1.93-fold), *gstp1a* (1.9-fold), *sod1* (1.59-fold), and *sod2* (1.42-fold) showed significant upregulation (p-value < 0.05) in males, but none of the oxidative stress genes showed significant upregulation in females. Notably, however, *sod2* was the only gene with a significant difference (p-value < 0.05) in the expression between males and females at 90 dpf.

## Discussion

Fluctuations in the temperature of aquatic environments can result in stress at cellular and physiological levels in aquatic animals (Podrabsky and Somero, 2004; Dong et al., 2006; Folkedal et al., 2012; Terrazas et al., 2017; Sherif et al., 2024); however, the long-term effects of thermal stressors on fish during early development are not well-known. In the present study, after treating fish with daily thermal fluctuations from 0 to 30 dpf, most of the *hsp* genes we measured exhibited significant upregulation in juvenile and adult fish, suggesting delayed cellular effects caused by the thermal stress during the larval stage. This increase might be because of the role of *hsp* genes in the reassembly of structures damaged by heat shock (Lindquist and Craig, 1988). In our study, *hsp90aa1, hsp90ab1*, and *hsp47* were upregulated in the juvenile and adult stages.

Heat shock protein 90 (HSP90) plays an important role in the final function of proteins (Li and Buchner, 2012), such as in the configuration of steroid hormone receptors (Prodromou, 2016). The amount and type of steroid receptors expressed in males and females are typically different (Voigt et al., 2009), which might partially explain the sex-biased expression of these proteins in adults in response to early-life temperature fluctuations in the present study. Heat shock protein 47 (HSP47) is involved in chaperone-assisted folding of collagen (Tasab et al., 2000). The upregulation of *hsp47* is also reported in injured tissue, for example, in fibrotic liver (Nagata, 2003). Thermal stress can increase oxidative stress and cause damage at the cellular level of tissues (Lushchak and Bagnyukova, 2006; Roychowdhury et al., 2021), which may partially explain the upregulation of *hsp47* in zebrafish.

The expression of genes involved in an oxidative stress response was also affected by temperature treatment during the embryonic and larval stages. The effect of temperature on oxidative stress genes is a phenomenon that has been reported in several species, such as *Onychostoma macrolepis*, *Acipenser fulvescens*, *Salvelinus fontinalis*, and *Sander lucioperca* (Yu et al., 2017; Bugg et al., 2020; Mackey et al., 2021; Chen et al., 2021). An upregulation of antioxidant proteins has been reported along with increased oxidation of biomolecules after exposure to higher temperatures (Musa et al., 2021). Antioxidant proteins can help cells detoxify free radicals and reactive species (Pamplona and Costantini, 2011). In our study, most oxidative stress response genes were upregulated at 60 and 90 dpf, especially in males. Male-biased oxidation of biomolecules, such as lipids, and increased expression of antioxidant proteins have been reported in adult *Neogobius melanostomus* exposed to thermal stress (Błońska et al., 2021), which is consistent with the male-biased expression patterns in our study.

The effects of early-life exposure to thermal fluctuations on the expression of genes included in our study were more pronounced in the juvenile and adult stages. Except for *glulb*, all the other genes quantified in this study did not show any significant change in expression during the larval stage due to the temperature treatment. Rather, most genes showed significant changes in expression in juvenile and adult fish. The similarity in the expression pattern of oxidative stress response genes in the temperature treatment group and control group might be because of higher expression of these proteins in the early-life stage of animals, a pattern that has been suggested to make individuals more resilient toward thermal fluctuations (Hsu et al., 2008; Almroth et al., 2010; Del Vesco et al., 2017). The expression of HSPs is known to change as animals age, and preconditioning animals with higher temperature in their early-life stage has been observed to affect the expression of these proteins in response to future thermal stressors (Tower, 2009; Hoffman et al., 2024). In addition, these proteins play an important role during early development and are expressed at high basal levels during early developmental stages (Krone et al., 1997; Loones et al., 1997; Krone et al., 2003). It is possible that natural higher basal expression of oxidative stress response and heat shock protein genes might also help with coping with harmful effects of thermal fluctuations, making the upregulation of these genes unnecessary. The upregulation of HSPs in the temperature treatment group in our study shows the delayed effects of exposure to higher temperatures at the cellular level, which also coincided with higher thermotolerance in adult fish.

Exposure to early-life thermal fluctuations significantly increased the CT_max_ of juvenile zebrafish in the present study, but this increase was not significant in adult fish. This increase in CT_max_ was detectable 30 days after the conclusion of the thermal fluctuations at 60 dpf, but it declined at 90 dpf. This suggests a delayed effect of early-life thermal experience on later thermotolerance that could be reversed by acclimation. The CT_max_ of fish has been shown to be plastic (Blair and Glover, 2019). Further, exposure to higher acclimation temperatures has been reported to increase the thermotolerance of species such as *Seriola lalandi, Salvelinus fontinalis, Acipenser fulvescens, Salvelinus confluentus, Oligocottus maculosus*, and *Danio rerio* (Fangue et al, 2011; Morrison et al., 2019; Bugg et al., 2020; Larios-Soriano et al., 2021; Best et al., 2025; Silva-Garay et al., 2025). Thermal preconditioning, in terms of repeated thermal shock or CT_max_, has not been reported to have a significant effect on the thermotolerance of some species, such as *Fundulus heteroclitus* and *Oncorhynchus mykiss* (Healy and Schulte, 2012; Guo et al., 2023).

However, experiencing thermal treatments such as repeated CT_max_, increased acclimation temperature, and previous thermal challenge has been reported to increase CT_max_ in *Danio rerio* (Morgan et al., 2018; Metz et al., 2025). Interestingly, Gavarikar and Craig (2025) did not observe a significant difference in the CT_max_ of fish that experienced diurnal thermal fluctuations during the larval stage after five months of acclimation to 28 °C, when compared to those raised at 28 °C their entire life. In our study, acclimating the zebrafish back to 28 °C significantly decreased the CT_max_ from 60 to 90 dpf, but there was no difference in CT_max_ between the temperature treatment and control group at 90 dpf. The difference in CT_max_ between our study and the Gavarikar and Craig (2025) results might be because in their study, the fish were acclimated to 28 °C for five months, compared to the 2-month acclimation to 28 °C in the present study. This longer acclimation time before conducting CT_max_ might be the reason for the difference in CT_max_ between our study and the results from Gavarikar and Craig (2025).

Experiencing different thermal conditions during the larval stage also affected the growth and size of the fish. Although no difference was found in the size of fish in the larval stage, fish that experienced thermal fluctuations were significantly smaller at the juvenile and adult stages. Exposure to high temperatures has been reported to increase metabolism in ectotherms (Resende et al., 2022; Aguilar et al., 2022). Although the relationship between metabolic rate and growth is complex and sometimes contradictory, there are studies showing increased metabolism along with decreased growth in other species exposed to high temperatures (Álvarez and Nicieza, 2004; Roychowdhury et al., 2019; Jourdain-Bonneau et al., 2023; Hinchcliffe et al., 2024).

Decreased growth, along with increased expression of cellular stress genes detected in our research, could be the result of increased metabolism in the temperature treatment group and the impacts of that remained even after ending thermal fluctuations.

In conclusion, our study shows the importance of early-life thermal treatments on cellular mechanisms, growth, and thermotolerance of zebrafish. Fishes have evolved adaptations for natural fluctuations in their environment, but changes in the thermal regime of ecosystems caused by anthropogenic activities can challenge their physiological limits. Our study shows that chronic thermal fluctuations during early life stages can increase thermotolerance measured via CT_max_ trials. However, this increase in thermotolerance coincides with a long-lasting increase in cellular stress and decreased growth. The effects of early-life temperature treatments on fishes may not be limited to the parameters measured in this study and may also involve other fitness-related traits, such as reproductive success and survival of animals, that should be investigated in future studies over the entire life cycle of the species. In addition, the difference in the expression of cellular stress response genes between males and females suggests the importance of sex-specific differences in aquatic animals for dealing with environmental challenges, which is worth further investigation.

## Acknowledgment

We appreciate the help and support provided by the Animal Care Staff at the Rady Biomedical Fish Facility and the Lindsey lab. This work was funded by the University of Manitoba Collaborative Research Program Grant and NSERC Discovery Grants to BWL and KMJ.

## Notes

### Competing Interest Statement

The authors have declared no competing interest.

